# Automated Seminal Root Angle Measurement with Corrective Annotation

**DOI:** 10.1101/2024.03.13.584803

**Authors:** Abraham George Smith, Marta Malinowska, Luc Janss, Anja Karine Ruud, Lene Krusell, Jens Due Jensen, Torben Asp

## Abstract

Measuring seminal root angle is an important aspect of root phenotyping, yet automated methods are lacking. We introduce SeminalRootAngle, a novel open-source automated method that measures seminal root angles from images. To ensure our method is flexible and user-friendly we build on an established corrective annotation training method for image segmentation. We tested SeminalRootAngle on a heterogeneous dataset of 662 spring barley rhizobox images, which presented challenges in terms of image clarity and root obstruction. Validation of our new automated pipeline against manual measurements yielded a Pearson correlation coefficient of 0.71. We also measure inter-annotator agreement, obtaining a Pearson correlation coefficient of 0.68, indicating that our new pipeline provides similar root angle measurement accuracy to manual approaches. We use our new SeminalRootAngle tool to identify SNPs significantly associated with angle and length, shedding light on the genetic basis of root architecture.

## 1 Introduction

Root architecture, encompassing root length, angles, distribution, branching pattern, and overall spatial arrangement, plays a pivotal role in resource capture and plant adaptation to varying environments [18]. The heterogeneous distribution of soil resources necessitates an efficient root system that can navigate gradients of nutrient and water availability [18, 28]. Seminal root angle has been identified as a valuable proxy for understanding the root system architecture of mature plants [5, 26].

A narrow root angle enhances water and nitrogen capture, whilst a wider angle facilitates foraging in the topsoil and improves phosphorus acquisition [20, 19]. The seminal roots are the initial root system, which determine the primary path for water utilisation in the early stages of plant growth. Traits such as seminal root angle, represent phenotypes suitable for diverse environments. A wider root angle tends to promote a shallow rooting system, ideal for areas with frequent in-season rainfall. In contrast, a narrow root angle associated with deep rooting improves access to deep soil moisture, which is particularly beneficial during dry growth periods [22, 8].

Domestication and selective breeding, primarily through indirect selection, shaped the root architectures observed in today’s cultivated varieties [30, 7, 10]. Investigating the genetic variation in different aspects of root architecture, including seminal root angle and root length, is key to understanding the underlying genetic factors controlling these traits. Marker-assisted selection strategies, reliant on specific quantitative trait loci (QTLs) associated with root architecture, represent a promising avenue for targeted root trait improvement.

Despite the establishment of several high-throughput methods for root phenotyping [32, 6, 38, 39, 34], obtaining measurements of traits such as seminal root angle remains challenging. Previous studies have relied on a manual process for root angle measurement [13, 16, 29, 17, 37], which is both time-consuming and prone to inter- and intra-annotator variation. Such manual processes have an effect on study design, limiting the scale of experiments and the frequency of imaging.

To the best of our knowledge, no prior work has addressed the need for an automated root angle measurement method. To fill this gap in the root phenotyping toolbox, we developed and evaluated an automated approach to measuring seminal root angle. Our approach utilises image segmentations created using interactive machine learning (IML).

We validated our pipeline using images of spring barley (*Hordeum vulgare*) roots from rhizoboxes, comparing automatic and manual angle measurements. To highlight the utility of our proposed method, we investigated the genetic architecture controlling root angle and length. This involved integrating root phenotypes obtained from our AI-based pipeline with genetic data by employing Bayesian variable selection, a robust analytical tool, to identify specific QTLs associated with root architecture [15].

To promote transparency and reproducibility, we make our image dataset freely available under a Creative Commons license at [this URL](https://zenodo.org/records/7870965#.ZEp5iXZByUk) and we open-source our code and make our downloadable installer available at [this URL](https://github.com/Abe404/SeminalRootAngle).

## 2 Materials and methods

### 2.1 Dataset

The spring barley population used in this study included 192 breeding lines and eight commercial cultivars (Cadiz, Skyway, Laureate, Pallas, Halfdan, RGT Planet, Flair, Prospect) bred for the Nordic climate. Plants were grown in custom-made plastic rhizoboxes (Figure 1; 20 cm x 2.5 cm x 37 cm) with a transparent plexiglass front plate. The rhizoboxes were inclined at 60° to encourage root growth along the visible side. Each box was filled with 1.8L substrate (a mix of turf, local topsoil and sand) and supplemented with 300 ml water. Four seeds of each line were sown in each box and thinned to two seeds per box three days after sowing (DAS). Two seeds per line were grown in three different boxes, providing six replicates per line. Due to a limited number of rhizoboxes and the screened population size, the experiment was conducted over four months (August to December), and the plants were phenotyped in 15 incomplete blocks of 44 genotypes. Cultivar Skyway was included in each block as a check line. The experiment was conducted in a single greenhouse compartment with additional heating and supplemental lighting (16/8 h and 21/18 ° C day/night). All lines in each block were allowed to grow for 12 DAS.

**Figure 1:**
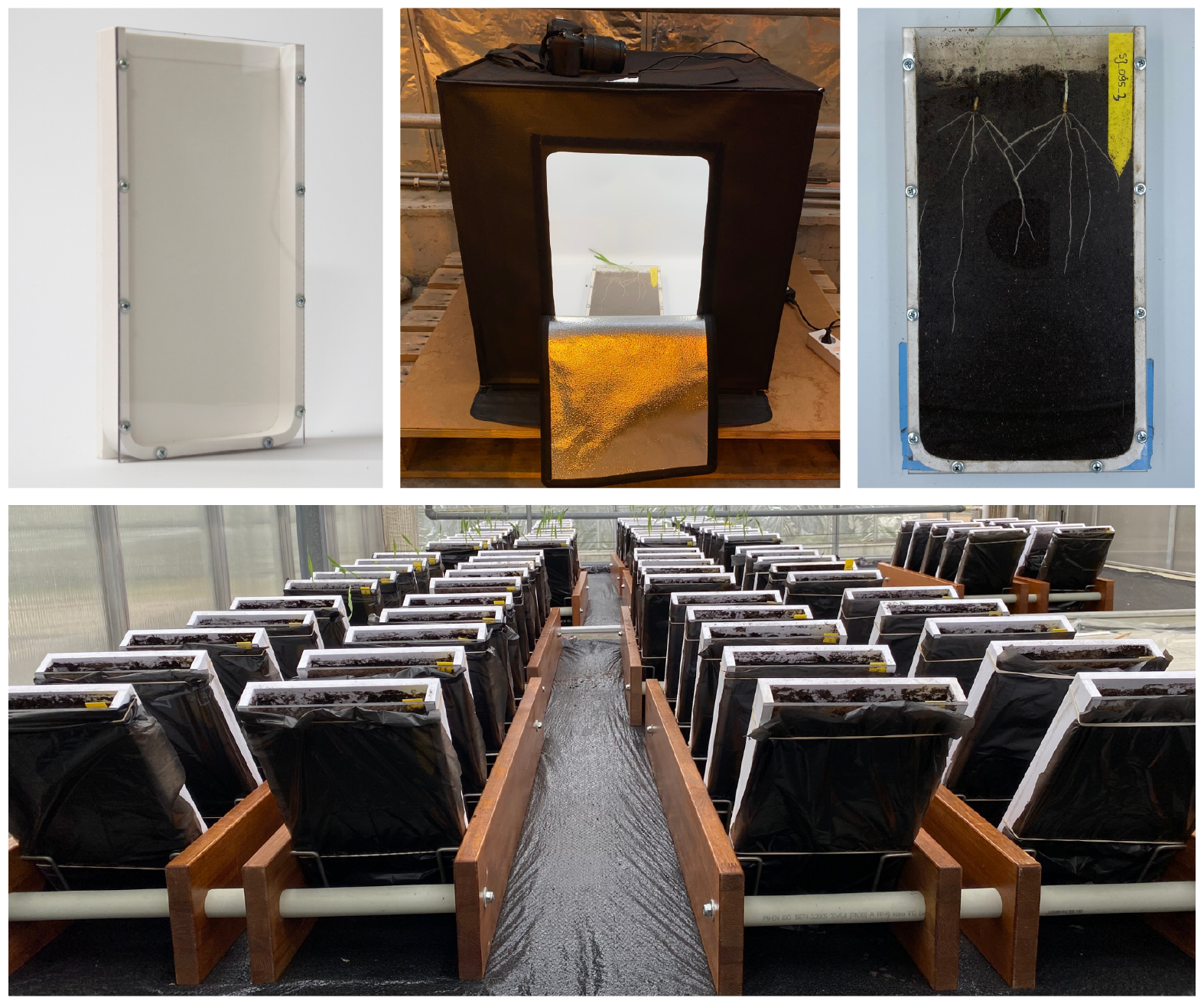
Spring barley plants growing in a rhizobox-based setup in a glasshouse. (top left) Empty rhizobox; (top centre) the light-box used in the experiment; (top right) an example of two plants in a rhizobox; (bottom) plants in the stands.

At 12 DAS, roots were imaged using a Nikon D7100 camera with a Nikon 18-140 mm 1:3.5-5.6 lens in a light-photo box (Figure 1). One high-resolution visual spectrum image (3050 x 5540 pixels) was taken of each rhizobox, and three pictures were taken of each line, resulting in 662 images in total, containing within them 1324 plants. Images were cropped and brightness corrected using a custom Python 3 script. Brightness correction employed the Yen thresholding method and intensity rescaling (supplementary material Listing 4.1)

### 2.2 Data split

To measure the generalisation of our root angle measurement method, we set aside a test set of 10% (n=66) of the images ^1^. The angle in these images was then manually measured using Fiji (Version 1.54). After excluding these 66 test images, 596 were available for segmentation model training.

### 2.3 Automated trait measurement

### 2.4 Overview of the automatic angle measurement algorithm

Our new automated root angle measurement method developed and evaluated in this study consists of two stages, firstly, using the rhizobox images (Figure 2a) a root segmentation (section 2.4.1) and seed point localisation segmentation (section 2.4.2) is obtained. Then the angle measurement process (section 2.4.3) combines segmentations to measure the angle of the primary roots from the images for each seed point present in the image.

**Figure 2:**
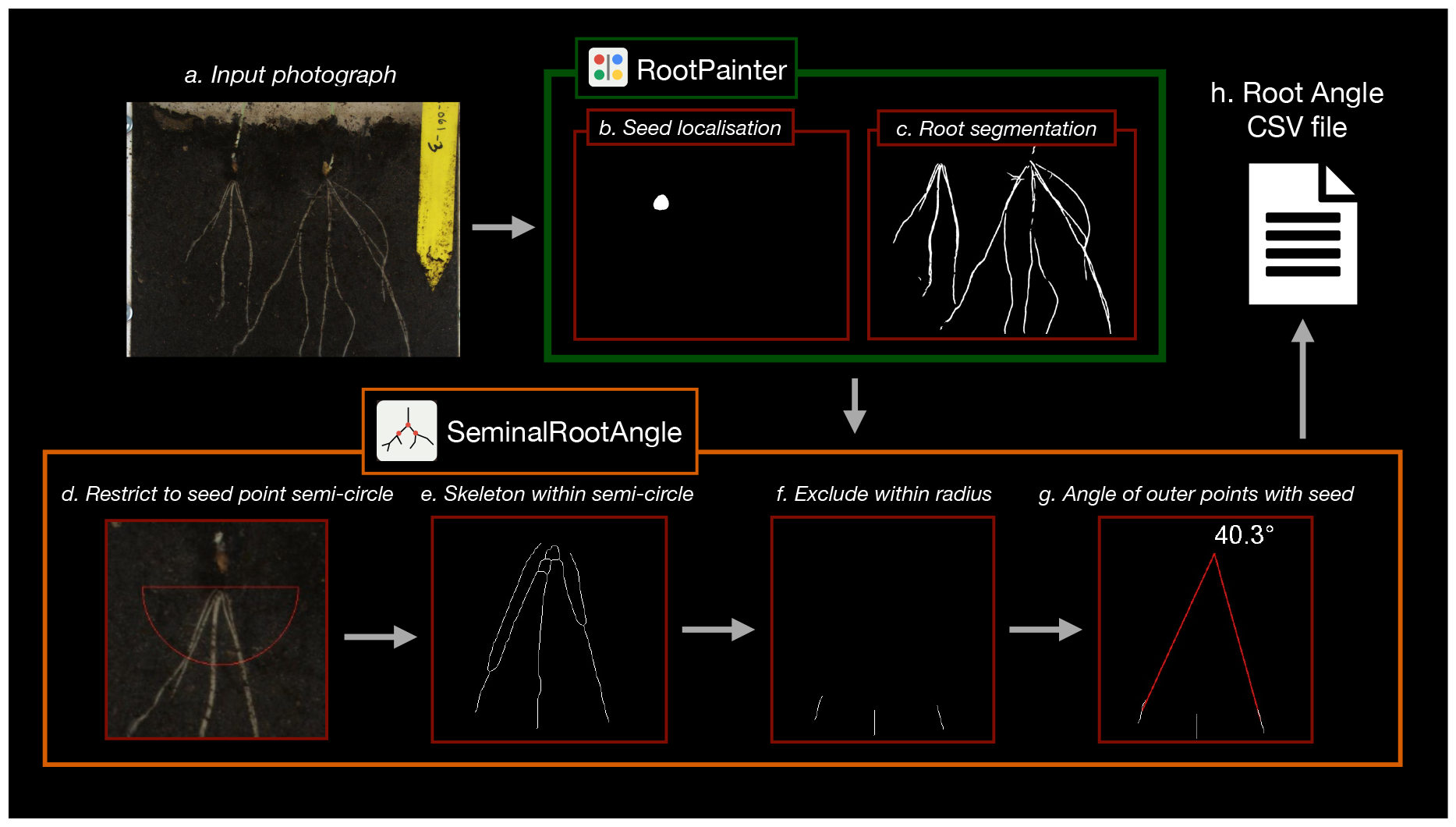
Root angle measurement pipeline. Including (a) input of a rhizobox photograph, which is segmented by two models for (b) seed localisation and (c) root segmentation. The diagram illustrates how these segmentations are then used in the newly developed SeminalRootAngle software, which (d) restricts to the seed point region, (e) skeletonises the segmented roots within this region, (f) excludes roots within a certain radius of the seed point and then (g) computes the angle between the outermost skeletonised roots, before (h) outputting to a CSV file including angles associated with all seed points found in all images.

The angle measurement algorithm, as implemented in our new SeminalRootAngle software, works by taking a region around the seed point (Figure 2d). The provided root segmentation is then skeletonised and restricted to this initial seed point region (Figure 2e). The region close to the seed point is excluded (Figure 2f), and the remaining outer pieces of the root skeleton are used to compute the angle (Figure 2e). The angles associated with all seed points are output to a CSV file (Figure 2h) including the image file name and their position in the image. In addition, to facilitate debugging, images showing each step of the processes for each seed point and a log of potential errors may be optionally output to a user-specified folder (Figure 4)

#### 2.4.1 Root segmentation

The root segmentation is used for calculating both the length and, in combination with the seed segmentation (section 2.4.2), the seminal root angle. To segment the roots, we created a root segmentation model using RootPainter. To obtain this root segmentation model, we first created a training dataset with a target size of 750 pixels width and height ^2^, sampling three tiles from each image.

We used the RootPainter corrective-annotation protocol, involving the annotation of clear regions of foreground and background in the initial images, and switched to corrective annotation [36], once the first model had been saved, which was detected by observing that the segmentation in the interface was beginning to approximate the structure of interest.

We started the network training procedure after the second image had been annotated and continued corrective annotation for a total annotation time of 3 hours. Once the time limit was reached, the final annotation was saved, even if only including partial correction of the errors in the image. We then left the model to finish training on its own on the annotations produced interactively. We also took a break halfway through annotation (after 1 hour and 30 minutes), where we let the model finish training. We used the contrast enhancement feature in RootPainter to enable darker regions of the images to be seen more clearly, as the lighting setup was sub-optimal in this study.

We then used an ensemble^3^ of the last five output model files to segment the full root dataset.

#### 2.4.2 Seed point localisation

Our seminal root angle measurement pipeline requires first locating the approximate seed point. To locate the seed point we trained a seed point segmentation model in a similar way to Section 2.4.1 but with the following alterations.

The seed point localisation image dataset does not require fine-detailed segmentation, thus having a larger field of view to locate the seed accurately is preferred. Instead of extracting smaller tiles (subregions), we downsized the dataset. We reduced the training dataset to 25% size ^4^ We then cropped to the top region of the image where the seed point was contained using Fiji, as cropping to only the relevant region typically improves segmentation accuracy [35]. We then trained a seed segmentation model by annotating the seed region in red (foreground) and background regions in green, whilst leaving areas close to the seed as undefined (not annotated) (Figure 3).

**Figure 3:**
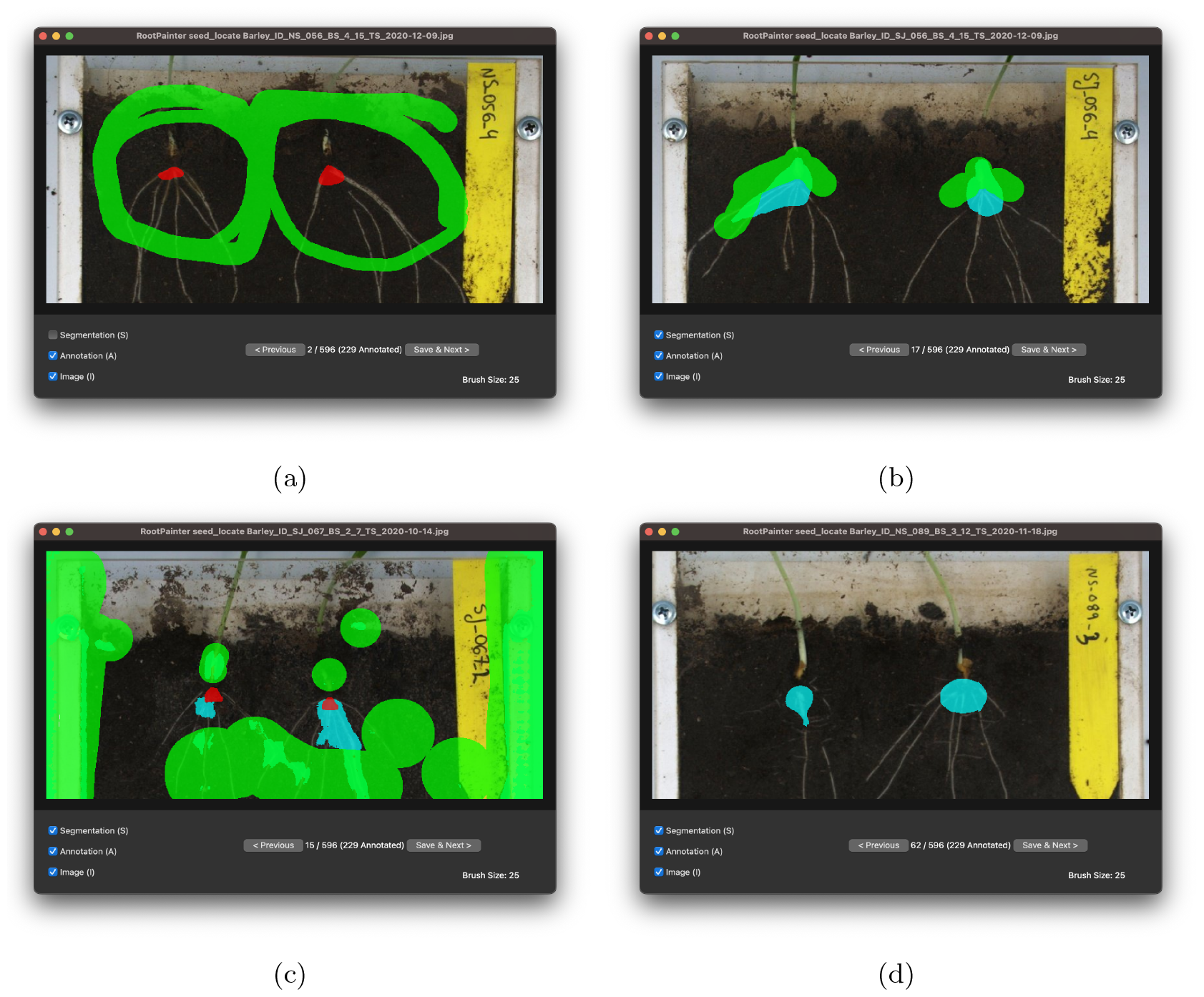
RootPainter interface showing corrective annotation at the various stages of model training for the seed localisation model. (a) Seed point location annotated as a triangle and background annotated in green further away. (b, c) Model starts to predict seed points with corrective annotation used to refine seed localisation segmentation model. (d) Seed point without correction.

Annotation was started by annotating the first clear image without regard to the model prediction (segmentation). We annotated a triangle for the seed location and annotated the background further away from the triangle as illustrated in Figure 3a.

The seed point foreground region did not correspond to a distinct physical object in the image but a region where the roots join the stem, which may be covered with soil as shown in Figure 3a.

After 15 images were annotated, the model started to detect the seed points. At this point, we proceeded with corrective annotation as shown in Figure 3b and 3c.

After progressing through 62 images, we encountered images that did not require correction, as shown in Figure 3d. We continued correcting mistakes in the segmentations for the entire training dataset. We spent less time training this model, with a total annotation time of 49 minutes and 15 seconds.

#### 2.4.3 Angle measurement

The root angle measurement function requires both the created seed point and root segmentations (Figure 4). Both segmentation folders must be specified as input parameters. An output location for the generated CSV and a folder for debug images, to allow results to be inspected, must also be specified. Measuring root angles from the spring barley image dataset containing 662 images with debug image output disabled, took 4 minutes and 30 seconds using a machine with an AMD Ryzen 9 5900X 12-Core Processor. We also output ‘debug images’ that allow inspection of the intermediary steps involved in the angle measurement process (described in section 2.4) and illustrated in Figure 2. When generating these debug images, the angle measurement process takes approximately twice as long.

**Figure 4:**
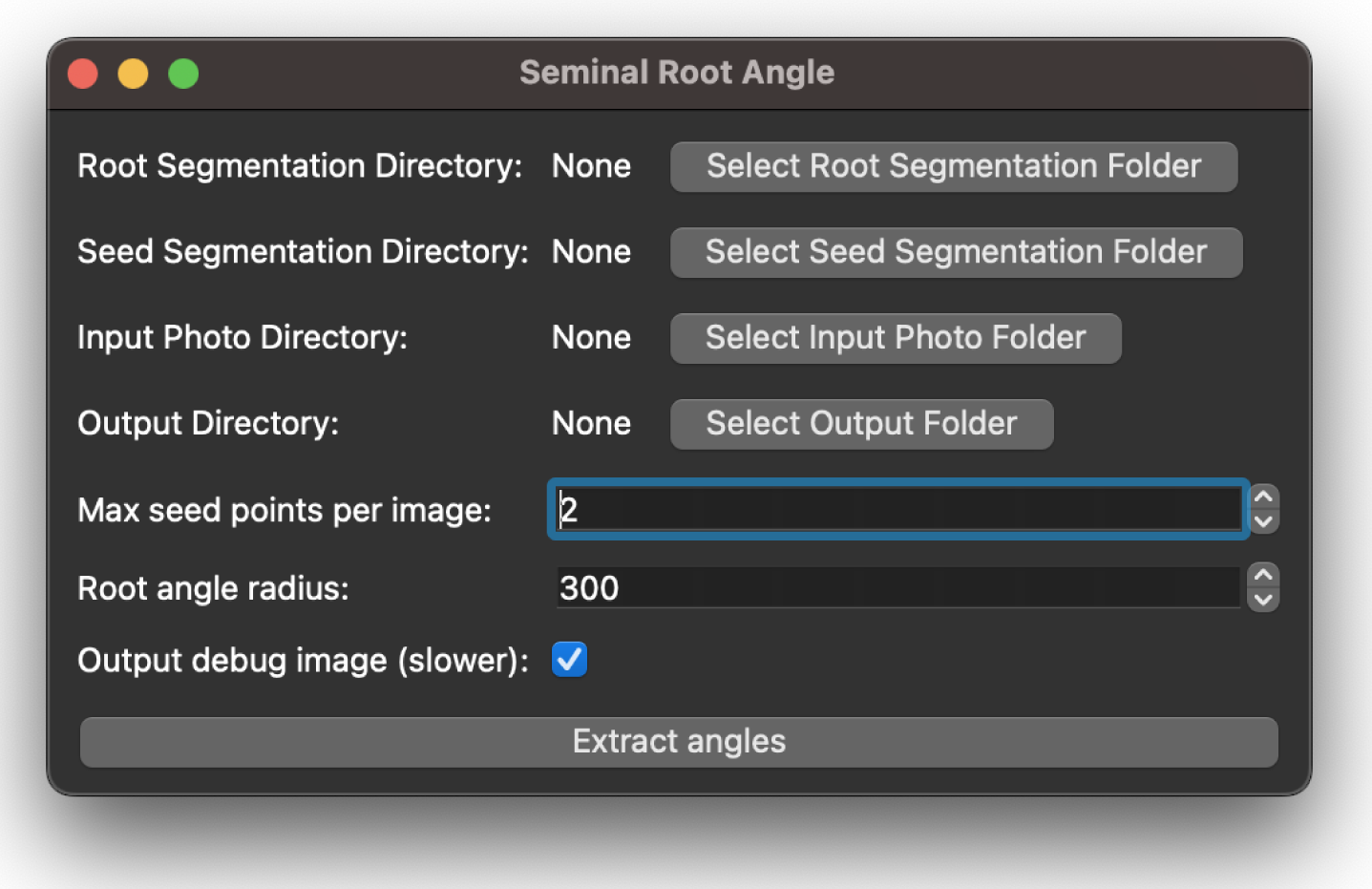
SeminalRootAngle GUI application

The SeminalRootAngle GUI also provides the option to specify the maximum number of seed points per image, which is useful for rhizobox images with multiple individual plants. A ‘Root angle radius’ can also be specified, which defines the local region in which to measure the angle around the seed point.

### 2.5 Evaluation of angle accuracy

To validate the automated angle measurements, we conducted manual measurements of root angles for 66 previously unseen images (corresponding to 132 seed points) in the test set. Manual angle measurements were conducted using Fiji’s dedicated ‘Angle Tool’, identified by a protractor icon. This tool allowed angle quantification by tracing lines representing the specific angles of interest within analysed images. To enable assessment of intra-annotator variability, three manual measurements were recorded for each image. To provide a comprehensive evaluation of the pipeline’s performance, automated measurements were compared to the average of the three manual measurements. We computed the Pearson correlation coefficient (r) and Mean Absolute Error (MAE) to assess the correlation strength and average absolute difference between the two measurement methods. To measure inter-annotator variation, 55 images (corresponding to 110 seed points) were measured by three different annotators. The inter-annotator analysis used a separate image set, distinct from the test-set used to compare manual and automated measurements. To provide insight into the efficiency of the automated angle measurement pipeline compared to manual methods, we measured the time required for conducting manual measurements.

### 2.6 Genomic association study

All individuals were genotyped using either the iSelect Illumina Infinium 4k, 9k arrays, or 15k SNP chip. Overlapping markers were then selected and used for the association study. To detect QTLs for root traits, we employed Bayesian Variable Selection (BVS) using a mixture prior on marker effects using the MCMC software BayZ^5^. The BVS approach used is based on George and McCullogh ([**george** ^**•**^ **1993**]) and developed for QTL mapping in Heuven and Janss ([14]); some details on the algorithms and implementation are given below. The fitted model for the total root length was as follows:

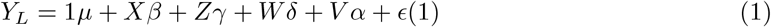

Where **Y**_*L*_ is the vector of total root length for *n* plants (replicates are not specified), *µ* is the model mean, **X** is a matrix with marker genotypes with dimension *n* × *p*, where *p* is the number of markers. The marker data was initially coded as -1, 0, or 1, and then column-centered to mean zero. The marker effects are represented by a vector ***β*** with a dimension of *p*× 1 and have a mixture distribution as detailed below.

**Z** is the design matrix capturing random effects for the block, ***γ*** is the vector of random effects parameters associated with the block, and 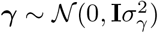. **W** is the design matrix capturing random effects for the position in the glasshouse nested within the block, ***δ*** is the vector of random effects parameters associated with the position in the glasshouse, and 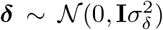. **V** is the design matrix capturing random effects for growth degree nested in ID, ***α*** is the vector of random effects parameters associated with growth degree days nested in ID, and 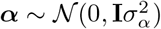.

Model residuals are 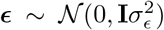. In the Bayesian setting, all model variance parameters 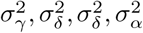, and 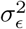 have prior distributions for which an (improper) unbounded uniform prior ∼ *U* (0, ∞) was used. In these distributions, *𝒩* () denotes a Normal distribution, and **I** an identity matrix of appropriate size.

The seminal root angle was modelled similarly to the total root length:

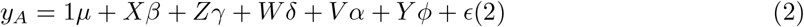

Where **y**_*A*_ is the vector of seminal root angles for *n* plants. Model (2) is mostly the same as (1), but an extra effect *Y ϕ* was added where **Y** is the design matrix capturing random effects for the combined effect of seed number, replicate, and block, 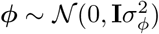 is the vector of associated random effects, and the variance 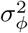 is treated the same as described above.

The BVS approach applied to marker effects ***β*** in (1) and (2) assumes a mixture of two normal distributions with a small variance for unselected/unimportant markers and a large variance for selected/large effect markers. An auxiliary indicator variable *ζ*_*i*_ ∈ {0, 1} for every marker effect *β*_*i*_ is used, so that 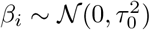 if *ζ*_*i*_ = 0 (small effects) and 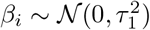 if *ζ*_*i*_ = 1 (large effects). Sparse variable selection is achieved by keeping 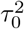 much smaller than 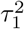 by constraining 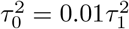, and by using Pr(*ζ*_*i*_ = 0) = *π*_0_, Pr(*ζ*_*i*_ = 1) = *π*_1_ = 1 − *π*_0_, where *π*_0_ is learned from the data with a prior distribution *π*_0_ ∼ Beta(100, 1) that implies *π*_0_ to be close to 1 and most marker effects to be small.

Implementation details include joint sampling of *β*_*i*_’s in blocks of 10 from conditional multivariate normal distributions, joint sampling of *ζ*_*i*_’s in groups of 2 after integrating out marker effects to improve mixing of the indicator variables, and the use of a Metropolis-Hastings sampler to update the variance parameters 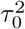 and 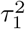 with the constraint fixed ratio.

The QTL mapping is based on using posterior means for the *ζ*_*i*_ indicator variable as a posterior probability of inclusion and a Bayes factor of the posterior and prior inclusion odds as a test statistic (see details in [14]). Computation of posterior means and standard deviations for all model parameters was performed using a single long Markov chain with a total of 250,000 iterations and a burn-in of 20,000 iterations.

## 3 Results

### 3.0.1 Validation of root angle

The mean automated root angle was 95.58 degrees, while the mean manual angle was 90.70 degrees. Measurements ranged from 43.20 to 151.30 degrees. There was a strong correlation between the manual and automated methods (r = 0.71) with a MAE of 12.9 degrees (Figure 5). Correlation coefficients between pairs of manual measurements from different annotators ranged from 0.61 to 0.80, with MAE values varying from 12.03 to 14.38 degrees (Supplementary Table 4.5). Coefficients between repeated manual angle measurements from the same annotator ranged from 0.82 to 0.9 (Supplementary Table 4.2).

**Figure 5:**
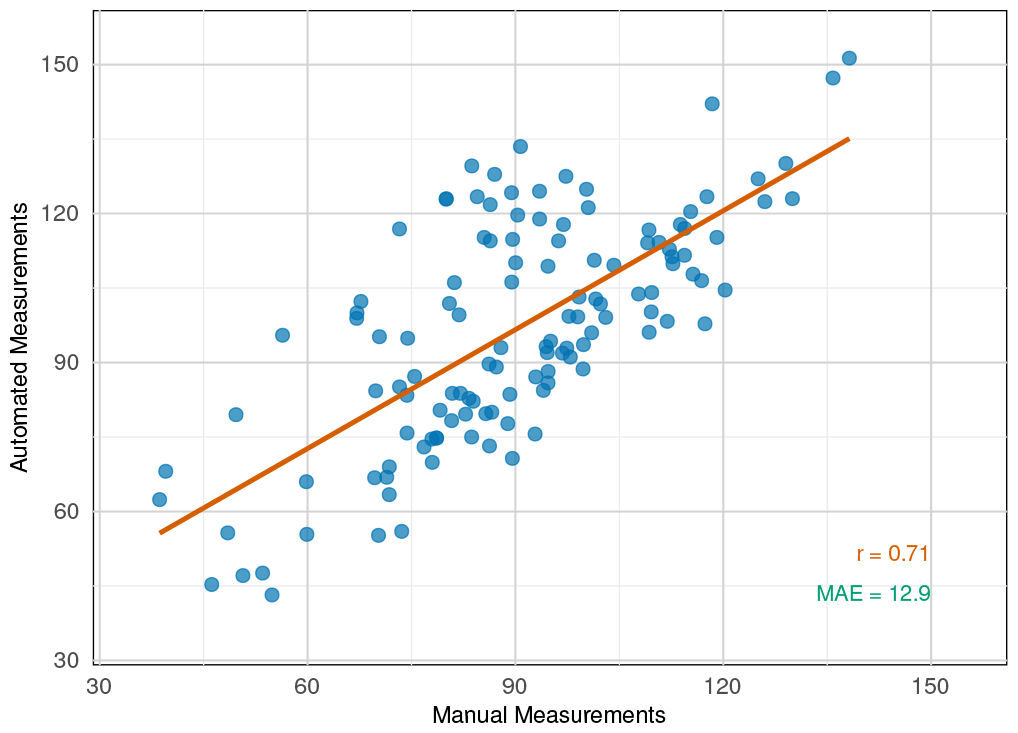
Scatter plot of manual vs. automated root angle measurements. The brick red line represents the linear regression line fitted to the data. r = Pearson correlation coefficient value; MAE = Mean Absolute Error.

On average, individuals performing manual measurements spent between 50 and 81 seconds on each image. None of the annotators did the measurements in one sitting; several breaks were necessary.

Images used in validation varied in quality, from high-quality with clearly visible roots and seeds to those with significant root obstructions and barely visible seed points.

We analysed the consistency and reliability of the correlation measurements across permutations of image order and increasing image counts (Supplementary Figure 4.3 and 4.4). In all cases, standard deviation approached zero, indicating minimal variability. Correlation coefficients stabilised consistently around the 100-measurements mark (Supplementary Figure 4.3 and 4.4), with slightly more iterations needed (around 120 measurements) for the correlation between automated and manual measurements. Fluctuations in standard deviation with increasing image count and varying order underscore the importance of selecting a sufficient number of images for validating the automated pipeline against manual measurements.

### 3.1 Bayesian variable selection

The Bayesian variable selection approach for TRL and SRA allowed us to identify significantly associated markers in the analysed population of spring barley (Figure 6 and Table 1). Two QTLs for TRL were identified, one within an intergenic region on chromosome 3H, and the other within two genes on chromosome 6H. The QTL for SRA was located within a gene body on chromosome 2H. A significant difference was observed for both traits in lines with a positive allele in all significant SNPs compared to lines with the negative allele (supplemental material 4.6).

**Table 1:**
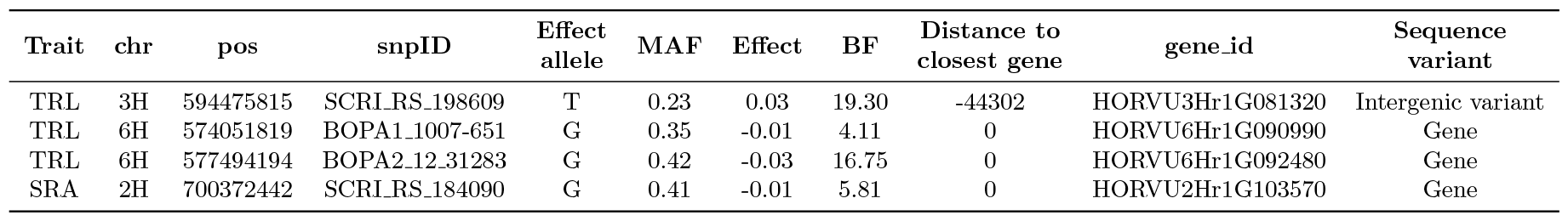
SNP from significantly associated genomic regions for total root length (TRL) and seminal root angle (SRA). MAF = minor allele frequency; Effect = allele effect of the SNP; BF = Bayes Factor; Where a value above 5 is considered as ‘strong’ evidence. SNP with BF between 3.2 and 5 are considered to be ‘putative.’

**Figure 6:**
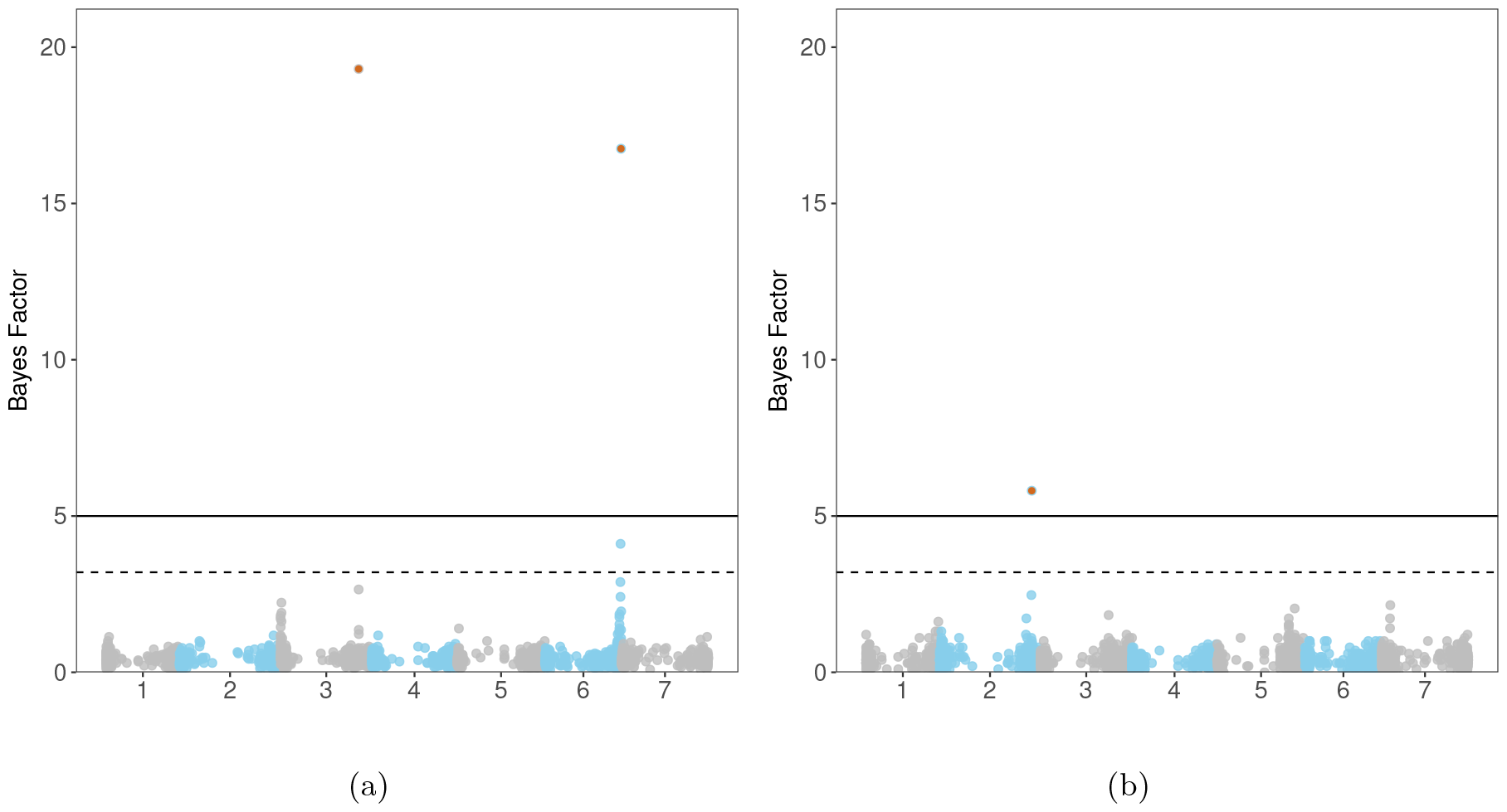
Manhattan plot of SNP for (a) the total root length and (b) seminal root angle. SNP above the line are considered significant; SNP above the dashed line are considered to be putative.

### 3.2 Computation requirements

The SeminalRootAngle GUI application is available with installers for Mac and Windows and can be run from source on Linux. The computational requirements are minimal. A computer with a dual-core CPU and at least 2GB of RAM should be sufficient, depending on the size of the images being processed. The RootPainter software used to generate the input segmentations is computationally demanding, with the server component requiring an NVIDIA GPU with at least 6GB of VRAM (We recommend at least 16GB). However, it is possible to run this via a freely available colab notebook. For further details, see [36].

## 4 Discussion

We present SeminalRootAngle, a high-throughput, user-friendly tool for measuring seminal root angles from rhizobox images. By speeding up root image processing, it proves useful for genetic studies, like Genome-Wide Association Studies.

Our proposed root angle measurement method offers speed, accuracy (depending on root visibility) and is freely available as open-source software with a user-accessible GUI. The segmentation component of our pipeline was implemented using RootPainter [34], which previous studies have demonstrated to be a flexible, accessible and efficient way to train segmentation models for a variety of root datasets [33, 3, 11, 12, 2, 21], including for images obtained from rhizobox experiments [1, 4].

The landscape of root image processing software has evolved significantly, with the emergence of several widely adopted, fully automated tools such as RootPainter [34], RhizoVision Explorer [31], WhinRhizo [38], and Digital Imaging of Root Traits (DIRT) [6]. Although RhizoVision Explorer provides the frequency of ‘shallow’, ‘medium’ and ‘steep’ root angles from the entire segmented root system [31], measuring seminal root or root spreading angles at the seedling stage has traditionally relied on manual methods [16] or semi-automatic software applications such as RootNav [23].

Despite the angle measurement accuracy supported by the strong Pearson correlation coefficient of 0.71 (Figure 5), there were occasional discrepancies between automated and manual measurements (Figure 5). These discrepancies highlight the inherent challenges of measuring root angles from images due to variability in root visibility caused by obstruction from soil debris and other objects.

The obstruction of roots poses a challenge for both automated and manual measurement, leading to inaccuracies and intra- and inter-annotator variation. Even with repeat measurements from the same annotator, the average Pearson correlation coefficient was only 0.85 (SD = 0.05). Likewise, when comparing measurements from three different annotators, we obtained a Pearson correlation coefficient of 0.68 (SD = 0.10), indicating substantial inter-annotator variability, which is likely due to the subjective nature of root angle measurement.

The impact of root visibility on accuracy is evident when comparing images with varying quality levels. Automated measurements closely align with manual measurements in high-quality images with clearly visible roots and well-defined seed points (Figure 7). However, significant root obstruction leads to deviations in automated measurements (Figure 7a). This occurs because the automated method relies on locating root segmentations within a defined region around the seed point, and soil covering the root in this area can hinder detection and result in inaccuracies. As root visibility decreases, both automated and manual methods encounter challenges in identifying the roots, thus affecting accuracy in the angle measurements (Figure 7).

**Figure 7:**
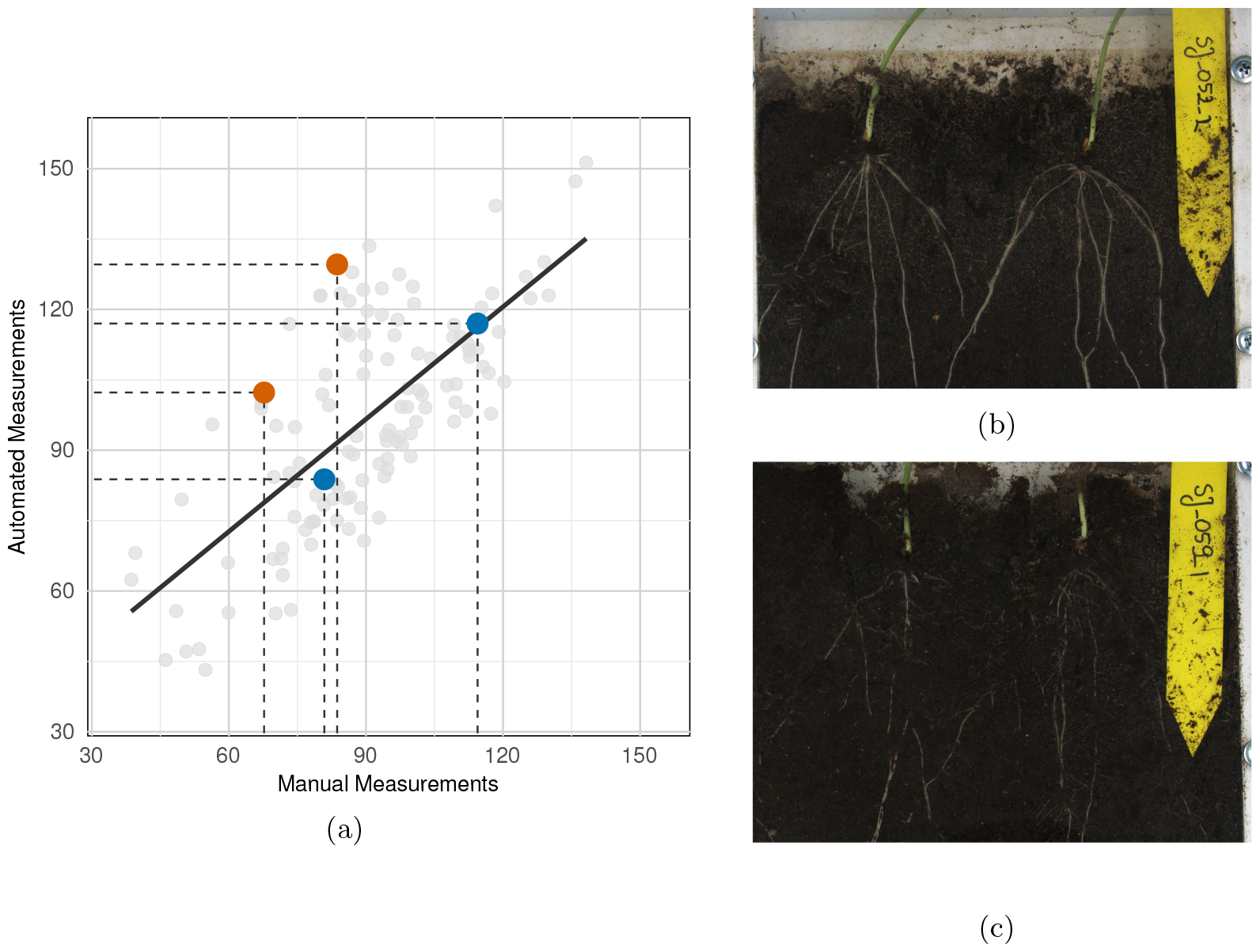
(a) Scatter plot comparing manual and automated root angle measurements, with highlighted points for two example images: (b) image with high root visibility (blue point) and (b) image with low root visibility (burnt orange points).

Despite facing the same root visibility challenges as manual methods, SeminalRootAngle achieves accuracy comparable to human annotators. Our analysis reveals that the automated method’s error margin falls within the range of observed variability in manual measurements, demonstrating its validity and reliability even in these challenging scenarios.

SeminalRootAngle presents a substantial improvement in efficiency compared to manual methods, particularly in terms of angle measurement time. Manual processing typically requires over a minute per image and can lead to annotator fatigue, necessitating breaks and potentially extending the process over days or even weeks for datasets of average size. In contrast, our proposed pipeline significantly shortens this timeframe. For instance, in our study involving 662 images, the average time to obtain two root angles (per one image) decreased to approximately 20 seconds, with a further reduction to only 4-5 seconds when focusing solely on angle measurement. Despite the initial time investment required for pipeline training, the efficiency gains become evident, especially with larger datasets.

We identified QTLs associated with seminal root angle and total root length, aligning with the broader goal of improving crop performance through enhanced root architecture [24, 19]. We found a single QTL linked to the seminal root angle located on chromosome 2H, consistent with previous findings by Jia et al. (2019) [16]. However, our results differed from Jia et al., who identified additional QTLs on chromosomes 3H, 4H, 5H, and 7H, and Robinson et al. (2018) [27] who reported a singular QTL on chromosome 5H. Additionally, we identified regions associated with total root length on chromosomes 3H and 6H. Farooqi et al. (2023) [9] recently reported 40 QTLs for nearly 16 root architecture traits, including two QTLs on chromosomes 2H and 7H for total root length. Our use of advanced lines from a breeding program, compared to studies using genetically diverse materials, might explain the lower number of identified QTLs, especially considering the higher marker density and diverse germplasm employed in those studies [16]. Nonetheless, our identified QTLs provide a foundation for allele stacking, offering breeders a strategic approach to optimise root architecture traits [25].

In conclusion, SeminalRootAngle stands as a valuable tool for root phenotyping, offering a swift, accurate and user-friendly solution for measuring seminal root angles from rhizobox images.

## Author contributions

A Smith implemented the root angle measurement pipeline and trained the models illustrated in the paper. M Malinowska performed the QTL analysis and evaluation of automated angle measurement accuracy. M Malinowska and A Smith wrote the manuscript. M Malinowska and AK Ruud planned and conducted the rhizobox experiment. L Janss wrote the genetic model. L Krusell and JD Jensen provided seeds for the experiment. T Asp obtained funding for the study. All authors read and provided comments on the manuscript

## Data availability

We make our image dataset freely available under a Creative Commons license at [this URL](https://zenodo.org/records/7870965#.ZEp5iXZByUk) and we open-source our code and make our downloadable installer available at [this URL](https://github.com/Abe404/SeminalRootAngle).

## Supplementary materials

**4.1 Brightness correction script**.

**4.2 Intra-annotator agreement**

**4.3 Inter-annotator correlation variation**

**4.4 Intra-annotator correlation variation**

**4.5 Inter-annotator correlation MAE**

**4.6 Root length and root angle QTLs**

## Footnotes

1 Using the random split method from the extras menu in RootPainter (Version 0.2.7).

2 Using the RootPainter ‘Create training dataset’ functionality.

3 Segmentation with ensembles is possible in RootPainter by selecting multiple model files when selecting the model to be used for the RootPainter ‘Segment folder’ function.

4 Using the resize method from the RootPainter extras menu.

5 http://bayz.biz/ and as R-package in https://github.com/ljanss/BayzR.

