## supplement for "Automated Seminal Root Angle Measurement with Corrective Annotation"

### Supplementary Material for paper “Automated Seminal Root Angle Measurement with Corrective Annotation”

#### 1 Brightness correction script.

Listing 1: Brightness correction of the rhizobox images

```
#!/usr/bin/env python3

"""
Generic Image Brightness Correction Script

This script applies brightness correction to a set of images using the Yen
    thresholding method.

Usage:
1. Place images in the input directory.
2. Run the script to process the images and save the corrected versions in
    the output directory.

Note: Make sure to install the required dependencies before running the
    script.

Dependencies:
- scikit-image
```

```
- imageio
- numpy
- os

"""
from skimage.filters import threshold_yen
from skimage.exposure import rescale_intensity
from skimage import io
import imageio
import os
import numpy as np

def main():
    input_directory = '/path/to/input_directory/'
    output_directory = '/path/to/output_directory/'

    # iterate through the names of contents of the folder
    for image_path in os.listdir(input_directory):

        # create the full input path and read the file
        input_path = os.path.join(input_directory, image_path)
        image_to_correct = io.imread(input_path)

        # correct brightness of the image
        yen_threshold = threshold_yen(image_to_correct)
        bright = rescale_intensity(image_to_correct, (0, yen_threshold), (0, 255))

        # Convert the image to uint8 format
        bright_uint8 = bright.astype(np.uint8)

        # Save the corrected image to the output directory
        output_path = os.path.join(output_directory, 'bright_'+image_path)
        imageio.imsave(output_path, bright_uint8, format='JPEG', quality=100)

if __name__ == '__main__':
    main()
```

#### 1.1 Intra-annotator agreement

| | M1 | M2 | M3 |
|---|---|---|---|
| M1 | 1.00 | 9.16 | 9.15 |
| M2 | 0.82 | 1.00 | 7.31 |
| M3 | 0.82 | 0.90 | 1.00 |

Table 1: Intra-annotator agreement and mean absolute error. Pearson correlation coefficients values between each pair are above diagonal, and Mean Absolute Error Values for each pair are below diagonal. M1:M3: refers to three manual measurements performed by the same annotator on the test set

#### 1.2 Intra-annotator Correlation Variation

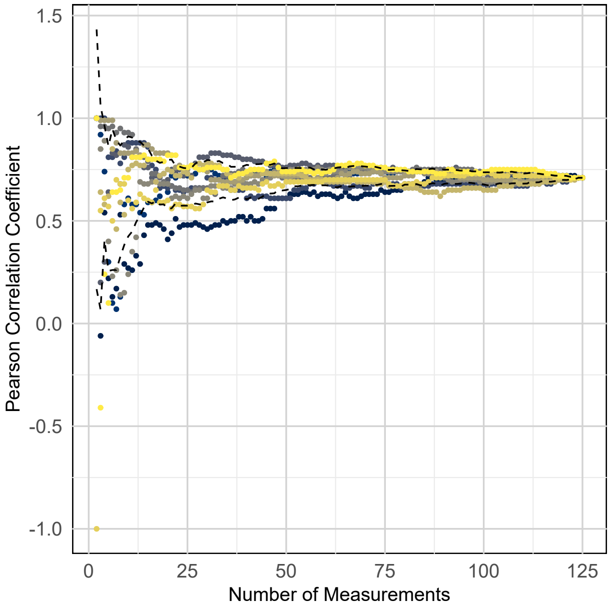


Figure 1: Variability of Pearson correlation coefficient across measurements for different permutations of image order. Points and colours indicate individual Pearson correlation coeffiecient measurements (correlations 1 to 10), Dashed line indicate standard deviation across all measurements.

#### 1.3 Inter-annotator Correlation Variation

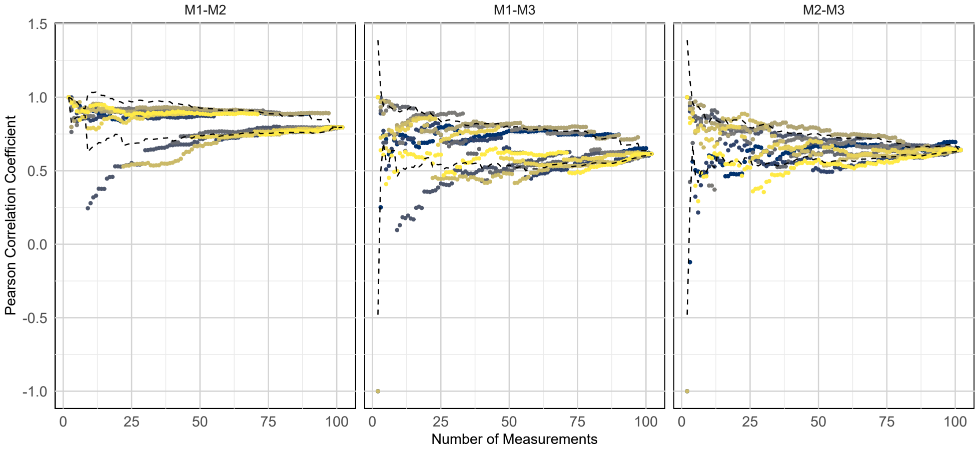


Figure 2: Variability of Pearson correlation coefficient across measurements for different permutations of image order between three different annotators. Points and colours indicate individual Pearson correlation coeffiecient measurements (correlations 1 to 10), Dashed line indicate standard deviation across all measurements.

#### 1.4 Inter-annotator Correlation MAE

| | M1 | M2 | M3 |
|---|---|---|---|
| M1 | 1.00 | 12.03 | 13.65 |
| M2 | 0.80 | 1.00 | 14.38 |
| M3 | 0.61 | 0.64 | 1.00 |

Table 2: Inter-annotator agreement and mean absolute error. Pearson correlation coefficients values between each pair are above diagonal, and Mean Absolute Error Values for each pair are below diagonal. M1:M3: refers to three manual measurements performed by three annotators on a set of 55 images

#### 1.5 Root length and root angle QTLs

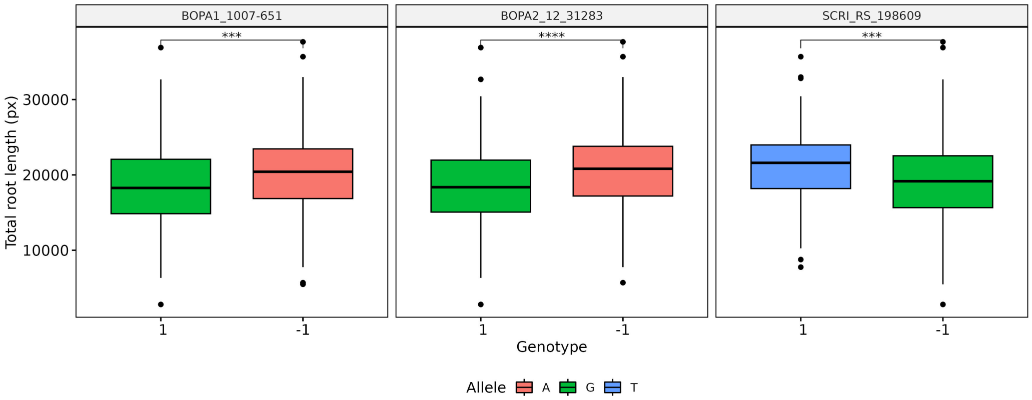


(a)

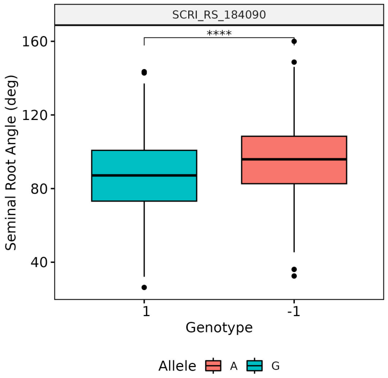


(b)

Figure 3: QTLs for root length and root angle traits.
